# Antibiotic resistance detection and concomitant species identification of ESKAPE pathogens by proteomics

**DOI:** 10.1101/2024.09.09.612008

**Authors:** Christian Blumenscheit, Yvonne Pfeifer, Guido Werner, Charlyn John, Franziska Layer-Nicolaou, Andy Schneider, Peter Lasch, Joerg Doellinger

## Abstract

Antimicrobial resistance (AMR) is an increasing challenge for therapy of bacterial infections. Currently, patient treatment is guided by antimicrobial susceptibility testing (AST) using phenotypic assays and species identification by MALDI-ToF biotyping. Bacterial phenotype prediction using omics technologies could offer several advantages over current diagnostic methods. It would allow species identification and AST to be combined in a single measurement, it would eliminate the need for secondary cultivation and could enable the prediction of phenotypes beyond AMR, such as virulence. In this study, the potential of proteomics for clinical microbiology was evaluated in an analysis of 126 clinical isolates covering 16 species including all ESKAPE genera and 29 of the most common AMR determinants. For this purpose, a flexible workflow was developed, which enables to report the AMR phenotype and the species of primary cultures within 2h. Proteomics provided high specificity (99.9%) and sensitivity (94.4 %) for AMR detection, while allowing species identification from very large sequence databases with high accuracy. The results show, that proteomics is well suited for phenotyping clinical bacterial isolates and has the potential to become a valuable diagnostic tool for clinical microbiology in the future.

## Introduction

Antimicrobial resistance (AMR) of bacteria is a major and emerging threat for public health. A systematic study of the global burden of AMR reports that 4.95 million deaths in 2019 were associated and 1.27 million deaths were attributed to resistant bacteria [1-4]. AMR is a major challenge for clinicians to treat bacterial infections. The selection of an appropriate antibiotic depends on the results of bacterial species identification and antibiotic susceptibility testing (AST) in clinical diagnostic laboratories [5, 6]. Nowadays, the taxonomic identity of clinical isolates is mostly analysed using MALDI-ToF MS-based biotyping, which is both rapid and cost-efficient [7]. On the other hand, AMR is usually analysed by phenotypic methods, which analyse the growth of bacteria in the presence of various antibiotics in parallel, such as broth dilution or disk diffusion techniques. Although phenotypic methods are well-established and powerful, genomics has gained a lot of attraction as a complementary method for predicting the AMR phenotype from the genotype [8, 9]. The use of molecular data for microbial diagnostics offers several potential advantages over phenotypic methods. It enables the integration of species identification and AMR detection into a single method, it omits the need for a secondary cultivation and offers the potential to predict the phenotype beyond AMR, e.g. the expression of virulence factors including toxins. Furthermore, those clinical data could directly be used for molecular epidemiology and so enable surveillance of bacterial infections in real-time. While an ISO-certified genomics workflow for identification and surveillance of antimicrobial resistance has been published recently, the potential of proteomics for clinical microbial diagnostics has not been investigated systematically, although the proof of concept has been demonstrated [10-12]. This is rather surprising as protein measurements are excellent predictors of the phenotype, which is why most clinically approved biomarkers and drug targets are proteins. Furthermore, antibiotic resistances are almost exclusively mediated by proteins and identification of bacterial species in clinical laboratories is already based on protein measurements [11, 13-15]. In addition, mass spectrometry-based proteomics has seen tremendous technological progress in recent years, which enables high throughput analysis of bacterial proteomes [16]. It has already been demonstrated, that thousands of proteins can be analysed in 1 min LC-MS/MS measurements and the depth of such ultra-high-throughput approaches is constantly increasing [17].

In this study, we aimed to establish and evaluate whole bacterial cell proteomics for antibiotic resistance detection and concomitant species identification of ESKAPE (*Enterococcus faecium, Staphylococcus aureus, Klebsiella pneumoniae, Acinetobacter baumannii, Pseudomonas aeruginosa*, and *Enterobacter* spp.) pathogens by analysing a representative panel of clinical isolates. ESKAPE pathogens are the most relevant agents of nosocomial infections and are closely linked with multidrug resistance and virulence. The sample panel consisted of 126 bacterial strains from 16 species, including all ESKAPE genera and 29 AMR determinants. The results demonstrate the enormous potential of proteomics for clinical microbiology, and the workflow utilized provides a solid foundation for further development of the technology in order to mature this type of analysis for clinical use.

## Methods

### Sample panel of ESKAPE pathogens

A total of 126 isolates from 11 bacterial genera spanning 16 species were included in this study (Table 1). This panel consists of the most prominent members of ESKAPE bacteria proposed by the WHO [5, 18, 19]. The extensively pre-characterized bacterial isolates were obtained from the strain collection of the Department of Nosocomial Pathogens and Antibiotic Resistance (FG13) of the RKI, which hosts the German National Reference Center for Staphylococci and Enterococci. A detailed list of the isolates and the typing of AMR and species prior to this study can be found in the Supplementary Appendix.

**Table 1:** Overview of the sample panel of multidrug-resistant bacteria including ESKAPE pathogens.

| Genus | Species | No. of Isolates | AMR Isoforms |
| --- | --- | --- | --- |
| *Acinetobacter* | *baumannii* | 15 | PER-7; AAC(6')-Ib-cr; OXA-23; OXA-80; OXA-72; PER-1; OXA-82; TEM-1; TEM; CTX-M-115; GES-11 |
| *Citrobacter* | *freundii* complex | 1 | AAC(6)-Ib-cr; CTX-M-15; NDM-1; OXA-1; TEM-1; OXA-9; GES-11; TEM; SHV-12; VIM-2 |
| *Enterococcus* | *faecium* | 25 | *vanB*; *vanSA*; *vanA*; *vanRA*; *vanRB*; *vanSB* |
| *Enterobacter* | *cloacae* complex | 2 | CTX-M-15; AAC(6); ACT-7; NDM-1; OXA-1; TEM-1; OXA-9; ACT; CTX-M-14; OXA-48; TEM; QnrA; VIM-4 |
| *Enterobacter* | *hormachei* | 2 | AAC(6')-Ib-cr; ACT; OXA-1; QnrA7; VIM-1; TEM; QnrA; SHV-12 |
| *Enterobacter* | *roggenkampii* | 1 | VIM-1; QnrS |
| *Escherichia* | *coli* | 19 | CMY-2; MCR-1; TEM-1; AAC(6')-Ib-cr; CTX-M-15; NDM-1; OXA-1; OXA-244; AAC(6')-like; OXA-2; QnrS1; QnrS; KPC-2; OXA-9; CTX-M-27; TEM; CTX-M-2; SHV-12; QnrB19; SHV-5; CTX-M; NDM-5; CMY-4 |
| *Klebsiella* | *grumontii* | 1 | AAC(6')-Ib; OXY-6-2; ACC-1; OXA-10; VIM-1; QnrS |
| *Klebsiella* | *oxytoca* | 1 | OXY-2-8; VIM-1; QnrS1 |
| *Klebsiella* | *pneumoniae* | 9 | CMY-4; KPC-2; NDM-1; SHV-1; AAC(6')-Ib-cr; CTX-M; OXA-232; TEM-1; SHV; AAC(6'); CTX-M-15; OXA-1; TEM; VIM-4; MCR-8.1; QnrS; SHV-5; VIM-1; SHV-11; CTX-M-14; CTX-M-27 |
| *Klebsiella* | *quasipneumoniae* | 1 | CTX-M-15; OKP-A-5; QnrS1 |
| *Morganella* | *morganii* | 2 | CTX-M-1; NDM-1; VIM-4; DHA-1 |
| *Proteus* | *mirabilis* | 3 | AAC(6')-Ib; DHA; OXA-10; VEB-5; CTX-M-15; AAC(6')-Ib-cr; DHA-1; NDM-1; OXA-1; TEM-1; ACC-1; KPC-2; QnrB2; QnrD |
| *Pseudomonas* | *aeruginosa* | 13 | PER-1; OXA-1; VIM-2; OXA-10; VEB-1; GIM-1; OXA-2; VIM-6; AAC(6'); IMP-7; IMP-15; VIM-28; IMP-16; IMP-1 |
| *Shigella* | *sonnei* | 3 | CTX-M-15; TEM-135; CTX-M-14; CTX-M-3 |
| *Staphylococcus* | *aureus* | 28 | *mecA*; *mecC* |

### Antimicrobial susceptibility testing (AST)

Antimicrobial susceptibility of all isolates was determined using broth microdilution (BMD) according EUCAST guidelines using interpretation standard EUCAST v14.0. Ten antibiotics were tested: ampicillin, cefotaxime, ceftazidime, cefoxitin, meropenem, gentamicin, kanamycin, nalidixic acid, ciprofloxacin and colistin. The AST of enterococci and staphylococci isolates was determined by BMD according to EUCAST standards and guidelines. For isolates without a clinical breakpoint, appropriate ECOFF values were used to distinguish the wild-type population from the non-wild-type (=resistant) population.

### PCR-based detection of resistance genes

The presence of various beta-lactamase genes was tested by PCR followed by Sanger sequencing for the majority of isolates using primers and protocols from previously published studies [20-22]. In addition, PCR screening and subsequent Sanger sequencing for genes contributing to resistance to aminoglycosides (*aac(6’)-Ib-like*) and colistin (*mcr-1*) were performed as recently described [23, 24]. The presence of vancomycin resistance gene clusters was determined by detecting the corresponding *vanA* and *vanB* ligase genes by multiplex PCR according to an accredited internal protocol [25]. Detection of methicillin resistance mediated by the genes *mecA* and *mecC* was determined by multiplex PCR according to an internal protocol [26].

### Next Generation Sequencing (NGS)-based detection of resistance genes

The publicly available Illumina sequence data were used to generate genomes. These short reads were filtered using *fastp* (v0.23.1, https://github.com/OpenGene/fastp) [27]. Genomes were generated using *unicycler* (v0.5.0, https://github.com/rrwick/Unicycler) with the *short-read-only* method using *spades* (v3.15.4) [28, 29]. Klebsiella genomes were screened for AMRs using *Kleborate* (v v2.2.0, https://github.com/katholt/Kleborate) [30]. For all other genomes, *abricate* (v1.0.0, https://github.com/tseemann/abricate) was used with the CARD database (v3.2.1).

### Bacterial cultivation and harvesting

All bacterial isolates were cultivated aerobically on Mueller Hinton II (Becton Dickinson, Heidelberg, Germany) agar plates at 37°C overnight. Bacterial cells were harvested by streaking with an inoculation loop and transferred to a 1.5 mL tube. The pellets were washed twice with 500 μL phosphate-buffered saline (PBS) and spun down at 6000 × g for 5 min.

### Sample preparation for proteomics

All samples were prepared using Sample Preparation by Easy Extraction and Digestion (SPEED) with minor modifications [31]. Samples were lysed in 100 μL trifluoroacetic acid (TFA) and mixed thoroughly with a pipette. The tubes were then incubated at 70°C for 3 minutes and subsequently neutralised by adding 1 mL of 2M Tris solution containing tris(2-carboxyethyl)phosphine (TCEP) to a final concentration of 10 mM and 2-chloroacetamide (CAA) to a final concentration of 40 mM. The tubes were incubated at 95°C for further 5 minutes. Protein concentrations were determined by turbidity measurements at 360 nm (1 AU = 0.67 μg/μL) using the NanoPhotometer® NP80 (Implen, Westlake Village, California).

Afterwards, 50 μL sample containing 50 μg proteins were mixed with 150 μL of rapid digestion buffer (Promega, Madison, Wisconsin, USA). Rapid trypsin (Promega) was added at a protein/enzyme ratio (w/w) of 10:1. Trypsin digestion was performed at 70°C for 15 min with shaking at 400 rpm in a thermomixer (Eppendorf, Hamburg, Germany). The peptide solution was acidified to a pH of ∼2 with 10 μL of 10% TFA and desalted using Pierce™ Peptide Desalting Spin Columns (Thermo Fisher Scientific) according to the manufacturer’s instructions. The desalted peptides were quantified by absorbance measurements at 280 nm using the NanoPhotometer® NP80 (Implen) and finally diluted with 0.1 % TFA to a concentration of 0.25 μg/μL.

### Liquid chromatography and mass spectrometry

Peptides were analysed on an EASY-nanoLC 1200 (Thermo Fisher Scientific) coupled online to a Q Exactive™ HF mass spectrometer (Thermo Fisher Scientific). 1 μg of peptides were separated on a PepSep column (15 cm length, 75 μm I.D., 1.5 μm C18 beads, PepSep, Marslev, Denmark) using a stepped 30 min gradient of 80% acetonitrile (solvent B) in 0.1% formic acid (solvent A) at 300 nL/min flow rate: 4–9% B in 2:17 min, 9-26% B in 18:28 min, 26–31% B in 3:04 min, 31–38% B in 2:41 min, 39–95% B in 0:10 min, 95% B for 2:20 min, 95–0% B in 0:10 min and 0% B for 0:50 min. Column temperature was kept at 50°C using a butterfly heater (Phoenix S&T, Chester, PA, USA). The Q Exactive™ HF was operated in a data-independent (DIA) manner. Full scan spectra were recorded in the m/z range of 345–1,650 with a resolution of 120,000 using an automatic gain control (AGC) target value of 3 × 10^6^ with a maximum injection time of 100 ms. The full scans were followed by 39 DIA scans with varying isolation window widths. DIA window placement was adjusted according to the retention time dependent distribution of peptide m/z values in 5 min intervals within the peptide elution window [16]. DIA spectra were recorded in the m/z range of 350–1,150 at a resolution of 30,000 using an AGC target value of 3 × 10^6^ with the maximum injection time set to auto and a first fixed mass of 200 Th. Normalized collision energy (NCE) was set to 27 % and default charge state was set to 3. Peptides were ionized using electrospray with a stainless-steel emitter (I.D. 30 μm (PepSep) at a spray voltage of 2.1 kV and a heated capillary temperature of 275°C. In order to minimize peptide carry-over, a wash run was included between each sample. Samples that still showed a peptide carry-over did not fulfil the required data quality criteria and were excluded from the study.

### Species identification based on LC-MS^1^ biotyping

Microbial species identification was performed as described previously, with minor modifications [32]. Briefly, peptide (MS^1^) feature lists were extracted from the DIA data. MicrobeMS (http://wiki.microbe-ms.com) was used to compare MS^1^ feature lists against a library of strain-specific *in silico* mass profiles obtained from UniProtKB/Swiss-Prot and Uni-ProtKB/TrEMBL protein sequence databases. Ranked lists of correlation or interspectral distance values (i.e. scores) were obtained, that provided information on the taxonomic identity of the organism under study (see [32] for details). The *in silico* database (v 2.0) consisted of 9031 strain-specific MS^1^ peptide mass profiles, each containing 8,000 – 15,000 different peptide mass entries. Variance-scaled Pearson’s product-moment correlation coefficients (Pareto scaling 0.25) were selected as distance values, using the mass region between 2000-5500 Da as input. The results of the correlation analysis were scored between 0 (no correlation) and 1000 (identity) and ranked to determine the taxonomic identity of the sample under study. In this way, identification of a sample by LC-MS^1^ biotyping was based on the identity of the top hit, which has the highest score in the score ranking list and therefore occupies the top (first) position in the list. The taxonomic identity of the isolates investigated throughout the study was determined beforehand using either MALDI-ToF MS, 16S rRNA sequencing, or whole genome sequencing (WGS). The accuracy of species identification was assessed by comparing the consensus taxonomic assignments of the reference methods and LC-MS^1^ biotyping. An identification was considered correct if LC-MS^1^ biotyping confirmed the taxonomic identity at the genus and species level. Cases counted as incorrect included misidentifications at the genus and/or species level and thus involved misidentifications of very closely related bacteria such as *Shigella sp*. and *E. coli* (see below). For the sake of simplicity, we used the accuracy of identification as a measure for overall identification quality. Identification test accuracy is defined as the ratio of the number of correct identifications to the sum of correct and incorrect identifications.

### Peptide identification

Peptides in the LC-MS/MS data were identified using DIA-NN (v1.8.1) [33]. Samples were analysed separately for each species in conjunction with the unrelated run option. Spectral libraries were predicted from the sample-specific species database and a pre-processed and reduced CARD database (March 2022) [34]. The sample-specific species database was selected according to the initial sample information. The respective reference proteome in the UniProtKB was used for library prediction and if no reference proteome was available, the proteome with the highest completeness (BUSCO value) was selected. The reduced CARD database consisted of the most common AMR determinants on the WHO watch list [3, 35], including 85 protein isoforms covering 74 AMR determinants (Table S 2 & Table S 3). Spectral libraries were predicted from the protein sequences using the deep learning algorithm implemented in DIA-NN with strict trypsin specificity (KR not P) allowing up to one missed cleavage site in the m/z range of 350 - 1,150 with charge states of 2 - 4 for all peptides consisting of 7-30 amino acids with activated N-terminal methionine excision and cysteine carbamidomethylation. Mass spectra were analysed with fixed mass tolerances of 10 ppm for MS^1^ and 20 ppm for MS^2^ spectra. The match between run (MBR) option was disabled. The FDR was set to 1 % for precursor identifications and the resulting precursor.tsv files were used for further processing in rawDIAtect.

### Antimicrobial resistance detection

Antimicrobial resistance was determined using rawDIAtect, which infers AMR from peptide identification results and summarizes the resistance phenotype of bacteria on various levels [10]. A minimum of 3 unique peptides was used to detect AMR determinants.

The results of proteomics-based antimicrobial resistance tests were analysed at three levels: AMR gene family, protein isoform and AMR phenotype. AMR status of the tested strains were previously characterized by PCR, Sanger sequencing and AST reporting. AMR detection by proteomics was considered true positive (TP) if the correct AMR gene family was detected. True negative (TN) test results corresponded to cases where proteomics confirmed the absence of AMR at the gene family, and protein isoform level. False negative (FN) or false positive (FP) AMR prediction corresponded to test results where the presence (FN) or absence (FP) of resistance could not be confirmed by proteomics.

Test parameters like the sensitivity and specificity were found to be helpful to describe the performance of the proteomics-based antimicrobial resistance test and were thus determined by means of the following well-known equations: sensitivity = TP / (TP + FN) and specificity = TN/(TN + FP), respectively. While the sensitivity refers to the ability of the test to detect a given AMR determinant, the specificity describes the test ability to confirm the absence of a given AMR determinant. For AMR protein isoform level and AMR phenotype level accuracy was calculated according to the equation: accuracy = correct / (correct + incorrect).

## Results

In this study, we aim to propose whole bacterial cell proteomics as an alternative method for antibiotic resistance detection and concomitant species identification of multidrug-resistant microorganisms including ESKAPE pathogens. Recently, we published a small-scale proof-of-concept (POC) proteomics study, which showed that simultaneous antibiotic resistance detection and species identification from DIA-MS data is possible with high specificity and sensitivity [10]. The diagnostic workflow was based on universal bacterial sample preparation using SPEED and data-independent acquisition mass spectrometry (DIA-MS), which potentially enables high-throughput proteomics but poses major challenges for data analysis, such as the identification of bacterial species from large peptide sequence data sets and the detection of low abundant AMR determinants. In the current study, we largely expanded the list of detectable AMR determinants and applied it to a sample cohort including all species and the most common resistance determinants of clinically-relevant ESKAPE pathogens among them *Enterococcus faecium, Staphylococcus aureus, Klebsiella pneumoniae, Acinetobacter baumannii, Pseudomonas aeruginosa*, and *Enterobacter* spp. A total of 126 isolates representing 16 bacterial species and 29 AMR gene families with 74 AMR isoforms were tested in this study (Table 1). Compared to our previous work, this study also includes significant improvements in the analytical workflow, which increased the throughput from 8 to 29 samples per day while maintaining a detection limit of < 10 protein copies per bacterial cell [16]. An overview of the proteomics workflow is shown in Figure 1.

**Figure 1:**
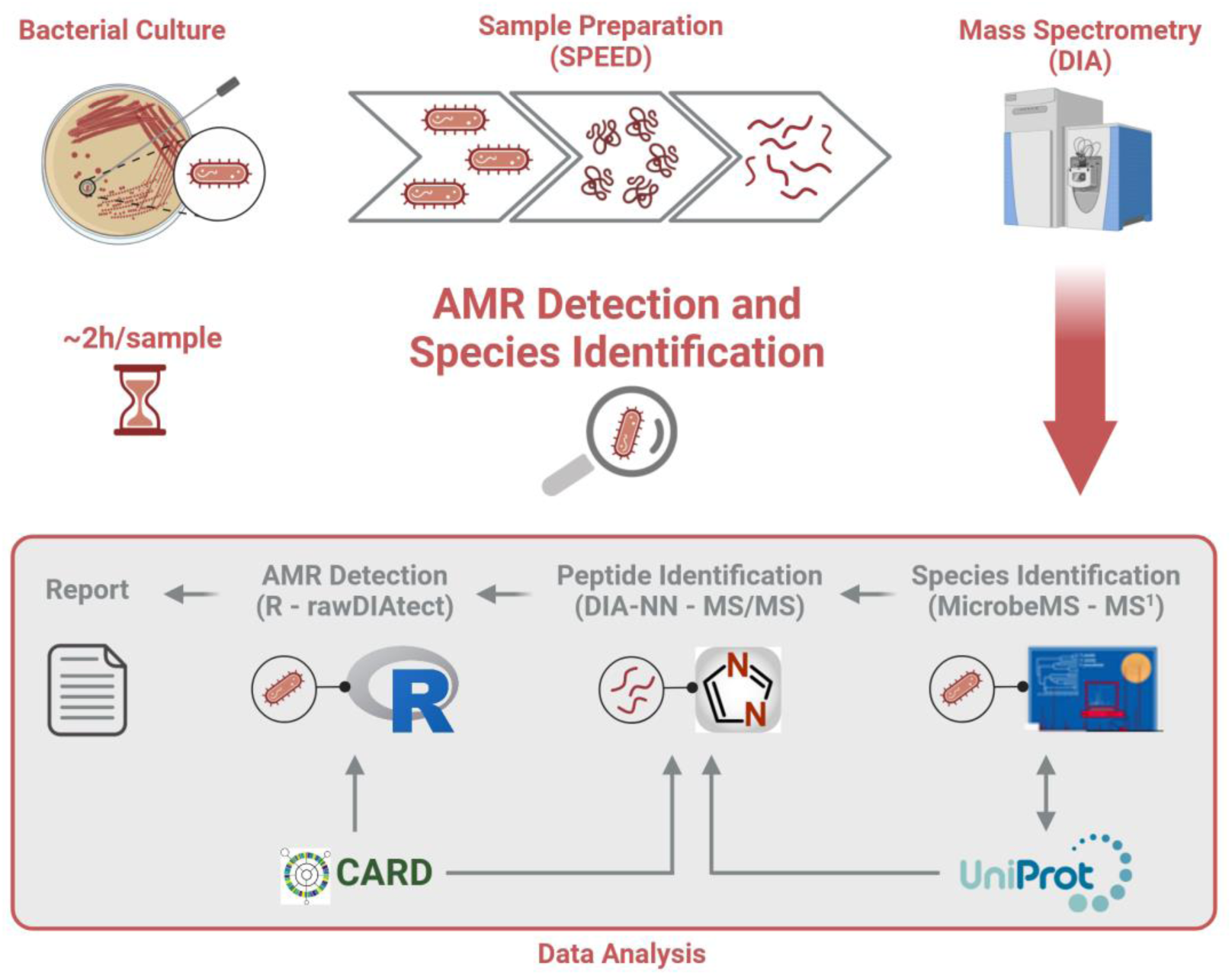
Proteomic workflow for species identification and AMR detection. Cultured bacteria are prepared for proteomics using sample preparation by easy extraction and digestion (SPEED), which is both rapid and universally applicable for all microorganisms. The resulting peptides are analysed using data-independent acquisition mass spectrometry (DIA-MS). Afterwards, the bacterial species is identified from the ^MS1^ spectra using MicrobeMS with a peptide mass barcode database build from the UniProtKB covering 9031 bacterial strains. The species information is used to predict a peptide spectral library for this specific sample, which is further enriched with sequences of AMR determinants obtained from the Comprehensive Antibiotic Resistance Database (CARD). This library is used to identify peptide sequences from the MS^2^ spectra using DIA-NN. The AMR phenotype is predicted from the identified peptide sequences using rawDIAtect, which summarizes the results on various levels, including AMR determinants, protein isoforms and drug classes. (Created in BioRender. Grossegesse, M. (2024) BioRender.com/r60u816)

### Species identification based on LC-MS^1^ biotyping

Bacterial species identification from DIA-MS data is a challenging task because peptide identification requires very large multi-species *in silico* predicted spectral libraries. Such libraries often contain more than 1 million protein entries, making analysis inefficient at best, if not impossible with most available software solutions. Therefore, we used an approach, previously published by us, that uses *in silico* generated peptide mass profiles and LC-MS^1^ data. The main idea of the MS^1^ approach is to generate an experimental mass list containing all MS^1^ peptide features of the microbial sample under study and compare it with sets of theoretically predicted tryptic peptide mass lists ultimately obtained from bacterial genome data. Correlations between the measured and the calculated peptide mass vectors are then determined and ranked so that the species identity can be inferred from the top-ranked entry.

This method is computationally intensive but allows rapid species identification from DIA-MS data. The results of the LC-MS^1^ biotyping analysis are summarized in Figure 2. In total, 117 out of the tested 126 species were correctly identified giving an overall accuracy of species identification of 91%. Perfect identification accuracy (100%) could be determined for all species except *Escherichia coli*, which was assigned as *Shigella sp*. in 12 samples, resulting in only ∼37 % accuracy of identification for *E. coli*. Given that *E. coli* and *Shigella* sp. are genetically closely related, if not indistinguishable, it was interesting to note that no *Shigella* isolate was misidentified as *E. coli*. The detailed list of all identifications can be found in supplementary Table S 10.

**Figure 2:**
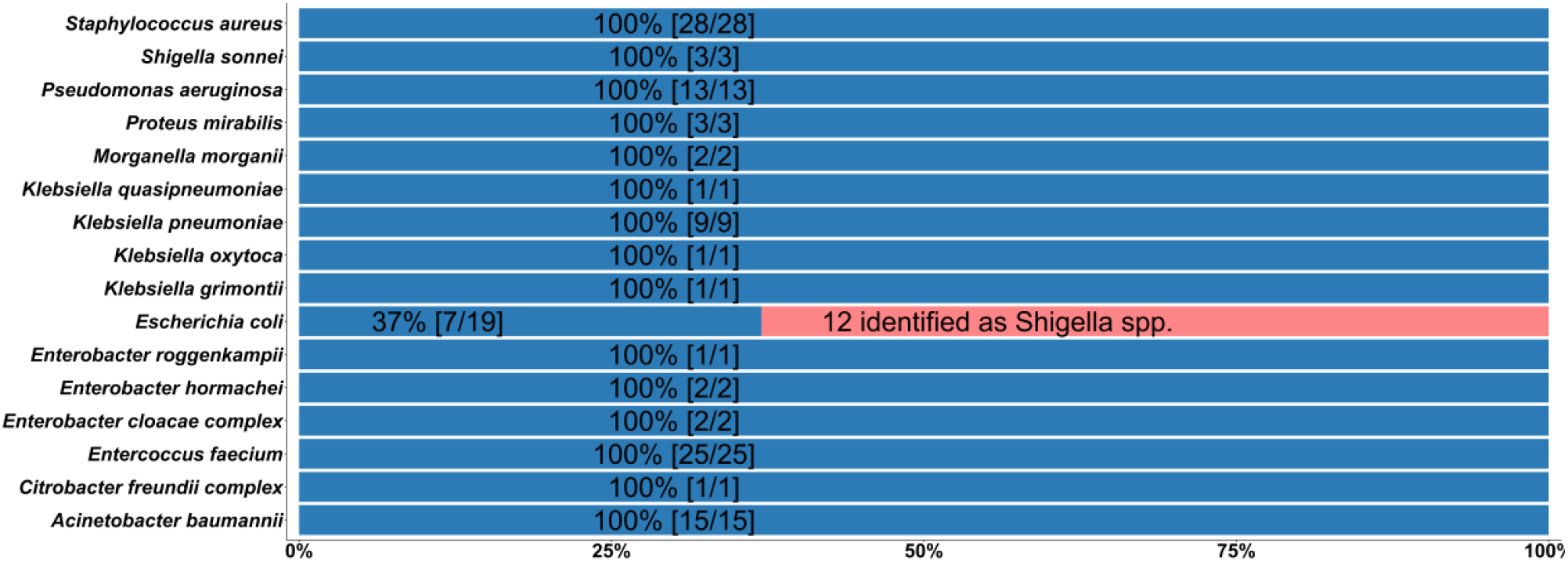
Species identification based on LC-MS^1^ biotyping. Distribution of identifications and resulting accuracy of species identification from LC-MS^1^ biotyping summarised by taxonomy. Overall accuracy was ∼91%. Total sample size and correct identifications are shown in parentheses. Twelve *Escherichia coli* strains were misidentified as different species of *Shigella*. For more information see supplementary table (*Table S 10* & *Table S 11*).

Accuracy of species identification for species other than *E. coli* in this case was comparable to or even higher than the accuracy of species identification reported for MALDI-ToF MS [7, 36-39]. It should be noted, that the *in silico* database used for the LC-MS^1^ biotyping approach has been obtained computationally and is therefore independent of measured spectral libraries, as in MALDI-ToF MS. Therefore, the database used for this study covers a large variety of bacterial species (9031 entries) and is thus comparable to available commercial MALDI-ToF MS databases.

### AMR detection by proteomics

In this study detection of antimicrobial resistance by proteomics relies on the protein expression of resistance determinants in the absence of antibiotics. It is currently not known, if all AMR gene families fulfil this prerequisite because for many determinants deep proteomic measurements are not available. The sample panel in this study covers 29 AMR gene families comprising 74 isoforms in 126 samples, which were initially characterized using AST, PCR/Sanger sequencing and/or NGS. In total, the proteomics results represented in this study are the equivalent of 3.654 PCRs – this test number would be required to detect the tested AMR determinants at the gene level in all samples by PCR.

Overall, AMR determinants were detected with a sensitivity of 94.4 % (272 TP and 16 FN test results) and a specificity of 99.9 % (3365 TN / 1 FP). AMR isoform detection accuracy of found AMR determinants equalled 83.8 % (227 correct and 44 incorrect), and the per sample phenogroup detection is 90.5 % (114 correct and 12 incorrect). The results demonstrate, that the proposed proteomic workflow achieved almost perfect specificity and high sensitivity in the analysis of ESKAPE pathogens (Figure 3).

**Figure 3:**
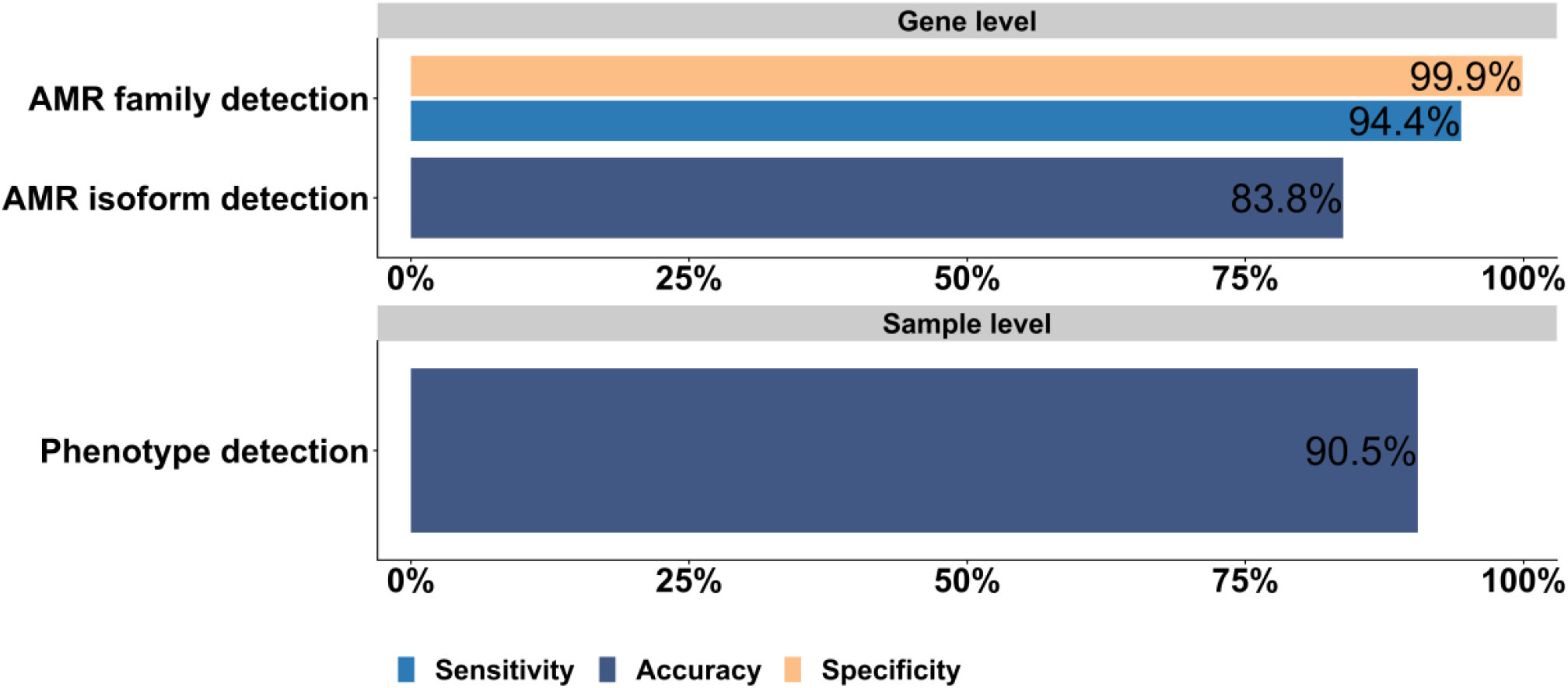
Performance of proteomics based AMR-diagnostics at the AMR gene family, AMR isoform and at the phenotype level. The sensitivity and specificity of AMR detection by proteomics in comparison to the methods phenotypic AST, PCR, NGS and Sanger sequencing is summarized for AMR determinants, protein isoforms and AMR phenotype. In total, 126 samples, containing 29 AMR gene families and 74 AMR isoforms were analysed. A detailed list of all classification results can be found in supplement Table S 1.

This is encouraging and demonstrates that the vast majority of proteins mediating a resistant phenotype are sufficiently expressed in the absence of antibiotics. However, the sensitivity varied between different AMR determinants. The proteomic results are summarized separately for each genus and AMR determinant in Figure 4. Overall, 21 of the 29 AMR determinants were detected with 100% sensitivity, while 5 AMR determinants were identified with sensitivities above 75%. The sensitivity for the detection of 3 AMR determinants was less than 75%. The low sensitivity of OXY was due to a single missed identification and is therefore probably resulting from the low incidence of this protein in the ESKAPE sample panel. In total, 8 out of the 16 missed AMR determinants were missed identifications of the quinolone resistance mediating protein Qnr, which could only be detected in 6 out of 14 Qnr positive samples (Sensitivity ∼43 %). Interestingly, most missing Qnr proteins were observed for the isoform QnrS1, which e.g. was not detected in all *Klebsiella* (0/4) samples. All AMR determinants could be found at least once in this study. A detailed list of all classification results can be found in supplemental Table S 1.

**Figure 4:**
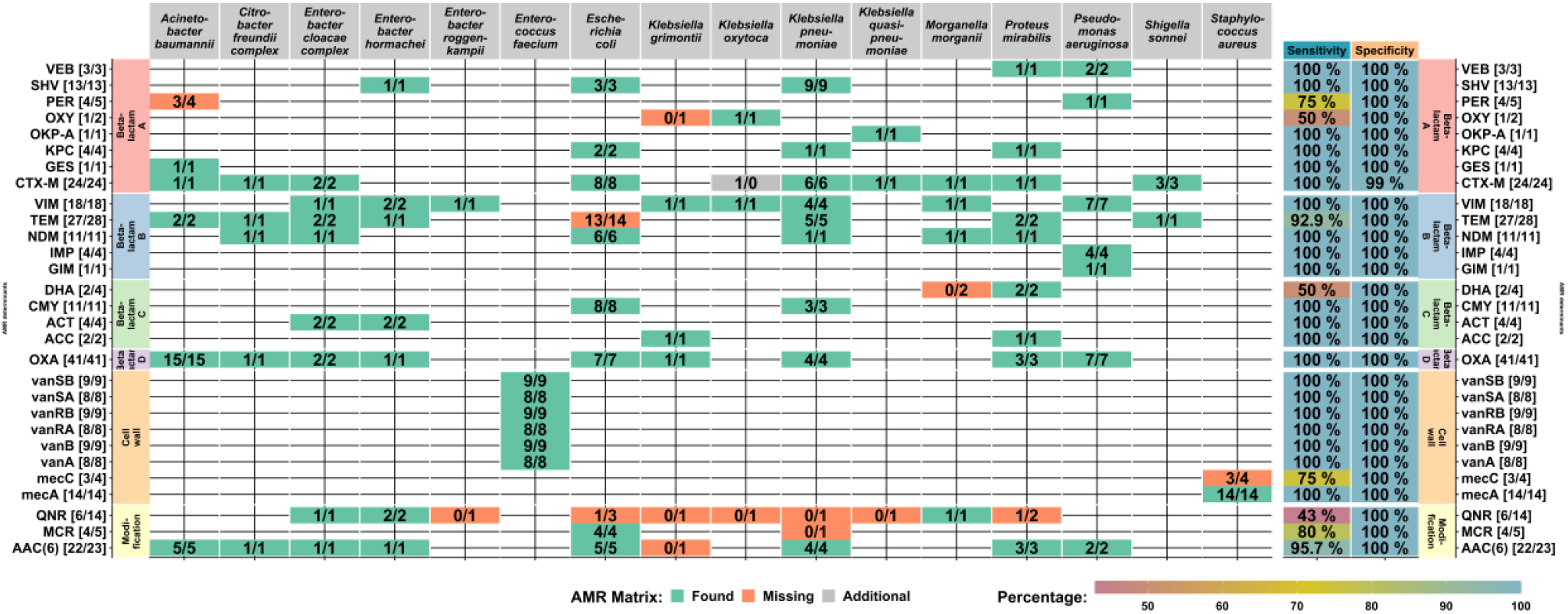
Results of the detection of AMR resistance determinants by DIA-MS proteomics sorted by AMR gene families (rows) and microbial species (columns) In this table coloured tiles encode the presence of AMR gene families for the given microbial species with green indicating accurate determination and red indicating partial, or complete missing confirmation of the AMR gene family. Numbers in the tiles denote the absolute number of AMR gene family detections by proteomics / the number of detections by the reference methods. The right columns colour code the sensitivity and specificity values of each gene family detection. A detailed list of all AMR characterization results can be found in supplemental table (Table S 5 & Table S 6 & Table S 7)

The resistance spectrum of each sample was further derived from the identified protein isoform and the results were grouped into corresponding phenogroups (Figure 5). Phenogroups are defined by the isoform and their corresponding resistance spectrum, a detailed list can be found in the supplement (Table S 8 & Table S 9). The correct resistance spectrum was determined in 114 samples out of 126 samples. The majority of incorrect classifications were due to the absence of quinolone resistance in 8 samples.

**Figure 5:**
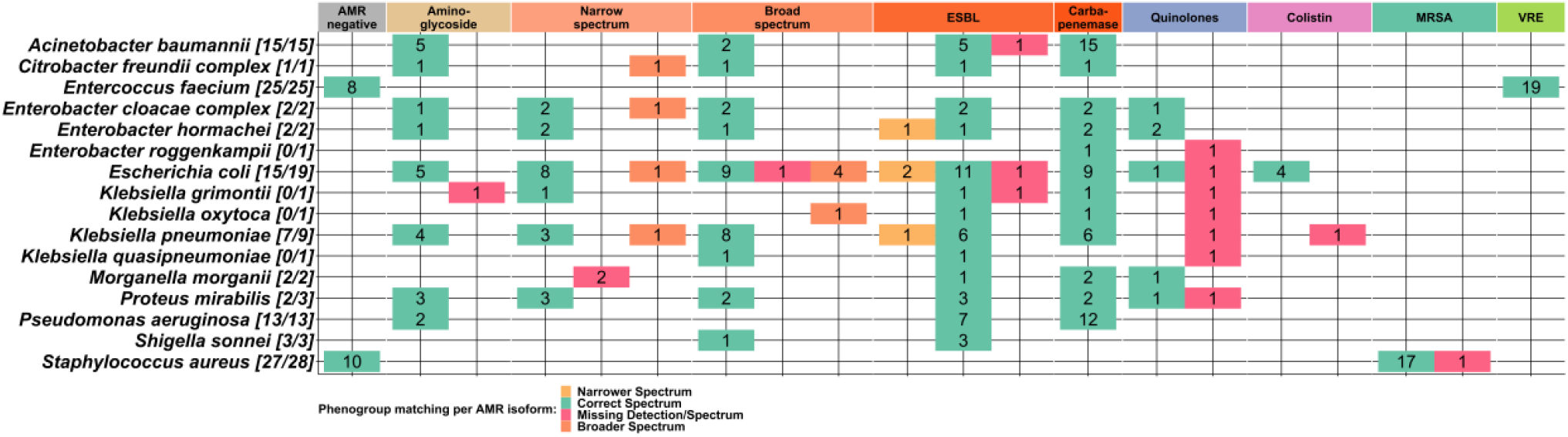
Detection of AMR phenogroups. The phenogroup detections based on AMR protein isoform identifications are summarized for each species. The tiles represent the phenogroup and the number of classifications. For each species the ratio of correct phenogroup classifications is shown in brackets. The sum of classifications per species can be higher than samples numbers due to multiple phenogroups per sample. The classifications are colour-coded. Tiles are green if the correct spectrum was found and red for a missing spectrum. If the detected phenogroup is overestimated (broader spectrum) the tile is coloured in orange and if the phenogroup was underestimated (narrower spectrum) the tile is coloured in yellow. For more information see supplement table (Table S 8 & Table S 9).

## Discussion

Molecular phenotyping of clinical bacterial isolates using omics technologies offers several advantages over current diagnostic methods. It would allow to perform both, species identification and antimicrobial susceptibility testing by a single measurement rendering secondary cultivation obsolete. Furthermore, omics technologies have the potential to predict bacterial phenotypes beyond AMR and e.g. enable the analysis of the expression of virulence factors including toxins and superantigens, which could improve patient care. Furthermore, such clinical data could directly be used for molecular epidemiology and so enable surveillance of bacterial infections in real time. While genomic methods are already well established and an ISO-certified workflow for AMR determination already exists, at least in Australia [40], there are no comparable proteomic methods. For a long time, this was due to serious technical limitations of LC-MS-based proteomics, which was only used in individual proof of concept studies for the targeted analysis of individual resistance determinants [13].However, the technical limitations in throughput, sensitivity and robustness of LC-MS have been overcome in recent years and the analysis of complete prokaryotic or low-complexity eukaryotic proteomes is now possible with a throughput of up to 180 samples per day [41]. As proteins shape the phenotype of cells, protein measurements should be very well suited to phenotype clinical bacterial isolates. In this study, we aimed to develop a flexible and universal workflow for bacterial proteomics and to determine the current potential of proteomics for characterizing ESKAPE pathogens within a sample panel of 126 clinical isolates. The results show that we were able to solve most of the technical problems for the successful implementation of proteomics in clinical microbiology. The acid-based sample preparation method (SPEED) is able to prepare all ESKAPE bacteria for measurement in less than 1h without protocol adaptations. The method also has great potential for automation to prepare bacteria for mass spectrometry without human intervention in the future. A DIA-MS setup subsequently allows the rapid analysis of bacterial proteomes with high throughput. In this study, data were recorded for 30 min for each sample, achieving a sensitivity of ∼ 4 protein copies per bacterial cell [16]. Initial studies show that these measurement types already allow thousands of proteins to be analysed in ∼ 1 minute of measurement time [17]. However, the complex structure of this data poses major challenges. The development of the LC-MS^1^ biotyping approach overcomes the limitation, that DIA-MS data cannot be analysed with very large sequence databases. The collapsing of the three-dimensional LC-MS/MS data into a one-dimensional peptide mass barcode enables to identify bacterial species within DIA-MS data even from databases derived from more than 9,000 genomes. The identification accuracy in this study was determined to be 91 % in total and 100 % excluding the discrimination issues between *E. coli* and *Shigella sp*., which are genetically very closely related and are difficult to discriminate even with currently established routine methods, such as 16S rRNA sequencing and MALDI-ToF MS. Therefore, LC-MS^1^ biotyping was found to already provide similar accuracy compared to the widely adopted MALDI-ToF MS biotyping method using spectral libraries predicted from genome sequence data, which roughly rivals the size of currently available MALDI-ToF MS databases. The species identification results enable to predict species-specific peptide spectral libraries and so enable the deep characterization of the cellular proteome. The most clinically-relevant AMR determinants were covered in the sample panel. The specificity of AMR determinant detection was 99.9 % in an analysis corresponding to 3654 individual tests, which shows that this approach is well suited for clinical diagnostics. Furthermore, all AMR determinants could be detected. This demonstrates, that proteins relevant for an AMR phenotype are expressed even in the absence of antibiotics, although their expression levels can be quite low. This fact was not known before as the majority of studies analysing gene expression in bacteria rely on transcriptomics. This is an important prerequisite for successful implementation of proteomics in phenotyping bacteria. The sensitivities of AMR detection were 94.4 % on determinant and 90.5 % on phenotype level. Those good values already underly the great potential of proteomics but also call for further improvements, as sensitivities were not evenly distributed among the different AMR determinants. Sample size for some determinants was too low for final evaluation, but especially the low detection sensitivity of 43% for quinolone resistance mediating protein Qnr needs further improvements. Interestingly, detection of QnrA and QnrB in *Klebsiella sp*. by mass spectrometry has been demonstrated with sensitivities of 80 % and 85 % using targeted assays before [42]. This shows that the detection of Qnr by proteomics is not limited in general.

The encouraging results presented in this study suggest, that proteomics is well suited for molecular phenotyping of clinical bacterial isolates. The proposed workflow can easily be integrated into commercially available LC-MS platforms with higher sample throughput, such as the timsTOF HT and the Orbitrap Astral, and will enable the analysis of 100 - 200 samples per day, which is likely to further increase in the future. In comparison to genomics the analysis of such high sample numbers is still very flexible as sample orders can be prioritized and cost and time scale linearly with sample numbers. For example, it would be inefficient to sequence few numbers of urgent samples on a high-throughput NGS platform and time-to sequencing result does not linearly decrease for low number of samples. In proteomics, the bacterial phenotype of e.g. an urgent sample from a sepsis patient could be analysed within ∼1 h from bacterial cell to report using a fast MS instrument. The ability to routinely analyse hundreds or even thousands of bacterial proteomes opens up questions on the potential benefits of proteomics for molecular surveillance. The integration of proteomics and genomics data might enable to better enlighten the mechanisms of emerging AMR phenotypes and identify and survey factors for certain virulence types. A high number of available bacterial proteomes with defined AMR phenotypes is also necessary to potentially build models to predict minimum inhibitory concentrations (MIC) of antibiotics and so directly guide personalized patient treatments.

## AUTHOR INFORMATION

## Author Contributions

The manuscript was written through contributions of all authors. All authors have given approval to the final version of the manuscript.

- J.D., A.S., C.B. and P.L. conceived the study and developed the method.
- C.B., J.D. and P.L. designed the experiments.
- C.B., C.J. and J.D. performed the proteomic experiments.
- C.B. and J.D. analysed the proteomic data.
- P.L. developed MicrobeMS and performed the LC-MS^1^ analyses
- C.B. developed the R package rawDIAtect.
- Y.P., F.L. and G.W. performed the characterization of the isolates (AST, PCR, Sanger sequencing).
- C.B., P.L, J.D., G.W. and Y.P. analysed the study data and wrote the manuscript.

## Access to proteomics data

The mass spectrometry proteomics data have been deposited to the ProteomeXchange Consortium (http://proteomecentral.proteomexchange.org) via the PRIDE partner repository with the dataset identifiers **XXXXXXX**.

## Access to code and programs

The R-code of rawDIAtect is available on https://github.com/CptChiler/rawDIAtect MicrobeMS can be downloaded from https://www.microbe-ms.com/

